# HUNTINGTON DISEASE ALTERS THE PATTERNING OF NEOCORTICAL AREA IN MICE

**DOI:** 10.64898/2026.05.12.724482

**Authors:** Camille Lafage, Leslie Ratié, Sandrine Humbert

## Abstract

**Background:** Huntington disease (HD) is a neurological disorder caused by an aberrant CAG expansion in the *HTT* gene, producing a mutant protein (mHTT). Although HD is classically characterized by adult-onset cortical and striatal degeneration, accumulating evidence suggests that altered cortical development may also contribute to disease pathogenesis.

**Objective:** We sought to investigate the impact of mHTT on neocortical patterning, which is a largely unexplored aspect of HD.

**Methods:** Using the HdhQ140 HD knock-in mouse model, we performed immunofluorescence and *in situ* hybridization to analyze the patterning of the cortex from embryonic day 10 to postnatal day 7.

**Results:** During embryogenesis, HTT expression exhibited a high medial-to-low lateral gradient in the neocortex, like that observed for key transcription factors involved in cortical patterning. Notably, HTT expression was absent from the cortical hem, a critical patterning center. In HD, the protein gradient remained unchanged whereas the expression in medial pallium seemed increased. During the early development of the cerebral hemispheres, the expression of morphogens and signaling pathways, including *Shh, Fgf8*, and *Wnt*/*BMP* genes, were disrupted in organizing centers, leading to altered expression of major neocortical transcription factors. At postnatal stages, the motor and somatosensory cortical areas were misplaced. These developmental alterations were associated with postnatal sensorimotor deficits relevant to HD.

**Conclusions:** Our findings demonstrate that HD-related neurodevelopmental alterations arise as early as embryonic day 10 in mice. This supports previous work suggesting that defects in brain development contribute to HD pathogenesis prior to clinical onset.

## INTRODUCTION

Huntington’s disease (HD) is a dominantly inherited neurological disorder that typically manifests during mid-to-late adulthood with psychiatric, cognitive and motor symptoms. It is characterized by progressive dysfunction and loss of striatal and cortical neurons. HD is caused by an abnormal expansion of a CAG trinucleotide repeat in the *HTT* gene encoding the huntingtin protein (HTT). The mutation is an abnormal expansion of a CAG nucleotide repeat in the *HTT* gene that encodes for an abnormal polyglutamine expansion in the HTT protein. While non-affected individuals carry 9-35 CAG repeats, HD patients harbor more than 36 repeats^**1**^.

Although HD is classically considered an adult-onset disease, we reported that cortical development is impaired decades before the appearance of the first symptoms. In HD mouse models, mutant HTT (mHTT) disrupts mitotic spindle orientation, interkinetic nuclear migration and cell cycle progression in neural progenitors, resulting in decreased cortical thickness during embryogenesis^**2**^. mHTT also interferes with neuronal migration and maturation^**3–6**^ and impairs axonal growth through microtubule bundling defects within the growth cone^**7**^. Suggestive as these data were, the field was lacking the evidence that HD affects human neurodevelopment. Recent neuroimaging studies revealed abnormal symmetry of the Sylvian fissure in HD mutation carriers, normally formed around the fourteenth gestational week, suggesting an embryonic origin of cortical defects^**8**^. Consistently, our analysis of human fetal tissue from mutation carriers demonstrated molecular and cellular defects in neural progenitors, including altered polarity, differentiation, ciliogenesis, and cell cycle regulation^**9**^.

Despite these advances, linking early developmental abnormalities to later HD pathology remains challenging. Notably, transient postnatal expression of mHTT or depletion of wild-type (WT) HTT is sufficient to induce adult HD-like pathology in mice, suggesting that developmental anomalies contribute to the evolution of the disease^**6**^. In line with this, we found that counteracting postnatal dysfunctions in HD mice when they first appear could prevent later the development of the disease^**10**^.

These data showed that not only brain development is altered in HD but that in principle, HD is preventable. Therefore, a better understanding of brain development in HD and its consequences for the pathology is essential for the design of future therapies that aim to correct developmental defects. In this study, we investigated whether neocortical patterning is affected in an HD mouse model.

## METHODS

### Study design

The HdhQ140 HD knock-in mouse model of HD was used in the study to elucidate differences in cortical patterning. We examined brains of embryos and pups from heterozygous Hdh^Q7/Q140^ and wild-type (Hdh^Q7/Q7^,WT) littermates obtained by crossing Hdh^Q7/Q140^ male mice with WT female mice. We used a combination of immunofluorescence staining, *in situ* hybridization labelling and behavioral tests to analyze cortical patterning alterations and consequences in HD.

### Animals

WT C57/Bl6J female mice were obtained from Charles River. Animals were maintained with access to food and water *ad libitum* and kept at constant temperature (19-22°C) and humidity (40-50%) on a 12/12 hour inverted light/dark cycle. All experimental procedures were performed in authorized establishments (Grenoble Institute of Neurosciences, license #B3851610008) in strict accordance with the local animal welfare committee (Comité Local Grenoble Institute Neurosciences, C1EA-04), EU guidelines (directive 2010/63/EU) and the French National Committee (2010/63) for care and use of laboratory animals under the supervision of authorized investigators (permission APAFIS#32445-2021062417002297 v3).

### Whole-mount and on slide in situ hybridization and immunofluorescence

Embryonic and postnatal brains (E10-P7) were dissected in PBS and fixed overnight at 4°C in 4% paraformaldehyde. Whole-mount embryos were dehydrated in methanol. For section analyses, tissues were cryoprotected in 20% sucrose, embedded in Tissue Tech Medium, and cryosectioned at 20µm in coronal or sagittal planes. *In situ* hybridization on sections and whole-mount tissues was performed using digoxigenin-labeled antisense riboprobes and alkaline phosphatase detection, as previously described^**11,12**^. The sequences of the probes used in this study are available in the referred articles: *Htt*^**10**^, *Shh*^**13**^, *Rorβ* and *Lmo4*^**14**^, *Cad8*^**15**^, *Emx2, Lhx2, Pax6, Bmp4, Sp8, Couptf1*^**11**^. *Fgf8* and *Wnt3a* were kindly gifted by A. Pierani and F. Causeret (Imagine Institute, Paris). Immunofluorescence was performed as described in Barnat *et al*. (2020)^**9**^, with cold acetone antigen retrieval for HTT detection. Mouse anti-HTT (CERBM, Cat#4C8; 1:500) and Alexa Fluor-conjugated secondary antibodies (Invitrogen; A11008; 1:500) were used.

### Images acquisition and analysis

Dorsal views of *in situ* hybridization of heads (E12.5) and brains (E18.5 and P7) were acquired with a Leica EZ4HD microscope associated to the Leica Application Suite EZ software. The hemisphere’s perimeters, anteroposterior and mediolateral width were measured using the ImageJ Fiji software. *In situ* hybridization images were acquired with a slide scanner Zeiss Axioscan Z1 and with a Macroscope and ANDOR Solution software for sections and E10.5 whole-mount *in situ* hybridization, respectively. Images of immunofluorescence of brain sections were acquired with a Zeiss LSM70 confocal microscope using Zeiss Zenblue software (Zeiss). Images were processed using Image J and QuPath softwares.

### Behavioral tests

In addition to the body weight monitoring of Hdh^Q7/Q140^ and WT littermates from P0 to P7, we performed a series of behavioral tests to assess primitive reflexes (grasping) and sensory-motor functions (righting and huddling). Our behavioral tests were adapted from those previously described in Braz *et al*. (2022)^**10**^ and Arakawa *et al*. (2015)^**16**^ and are shortly described below. Tests were completed on 9-15 litters (average litter size: 9; range: 5 to 11). All tests were video recorded, with an Olympus Camera fixed on a tripod.

#### Grasping

The appearance of grasp reflex in forepaws was assessed by the ability of the pup to grasp a thin rod stroking the palmar surface of each forepaw. The pups were maintained softly by the experimenter who touched and stroked against the palmar surface of a forepaw with a cone-tip and the responses were recorded. We evaluated the reflex by the absence (0) or the presence (1) of grasp reaction.

#### Righting reflex

One pup at a time from the litter is placed on its back on a horizontal surface for 10 seconds. The time it takes to turn itself on its ventral face is measured. Measures are averaged for each stage, genotype and sex.

#### Huddling

Two pups from the same litter and of the same sex, were isolated from the mother and put facing each other (∼3cm apart) in a 10×10cm arena. Free moving pups were recorded for 5 minutes. We analyzed the latency to form pairs, the number of interactions and their duration.

### Statistical analysis

For each experiment, animals from at least 3 independent litters were used. GraphPad Prism software (GraphPad Software, Inc.) was used for statistical analyses. Outliers were identified by using the ROUT method with Q=1% and removed for following statistical analysis. Normality of data distribution was assessed using the d’Agostino-Pearson omnibus normality test or the Shapiro-Wilk normality test, depending on the number of experiments. Appropriate parametric or non-parametric tests, as well as two-way ANOVA with Sidak’s post hoc test, were applied.

## RESULTS

### HTT expression is modified in HD in the developing mouse cortex

We examined the expression of HTT in WT and heterozygous knock-in HD mice in which the first exon of the mouse gene was replaced by the human exon 1 of *HTT* carrying 140 CAG repeats (Hdh^Q7/Q140^)^**17**^. At E12.5 in control brains, HTT displayed an expression pattern with a high medial to low lateral gradient within the progenitor cells of the neocortex, which was conserved at the rostral, middle and caudal levels (Fig. 1 A). In WT animals, HTT expression was absent from the cortical hem, located in the medial pallium between the cortical hem boundary (CHB) and the choroid plexus (Cp) (Fig. 1A). In HD samples, HTT expression level in medial pallium was increased (Fig. 1A, see white stars) and the exclusion area was less well defined at the CHB.

**Figure 1.**
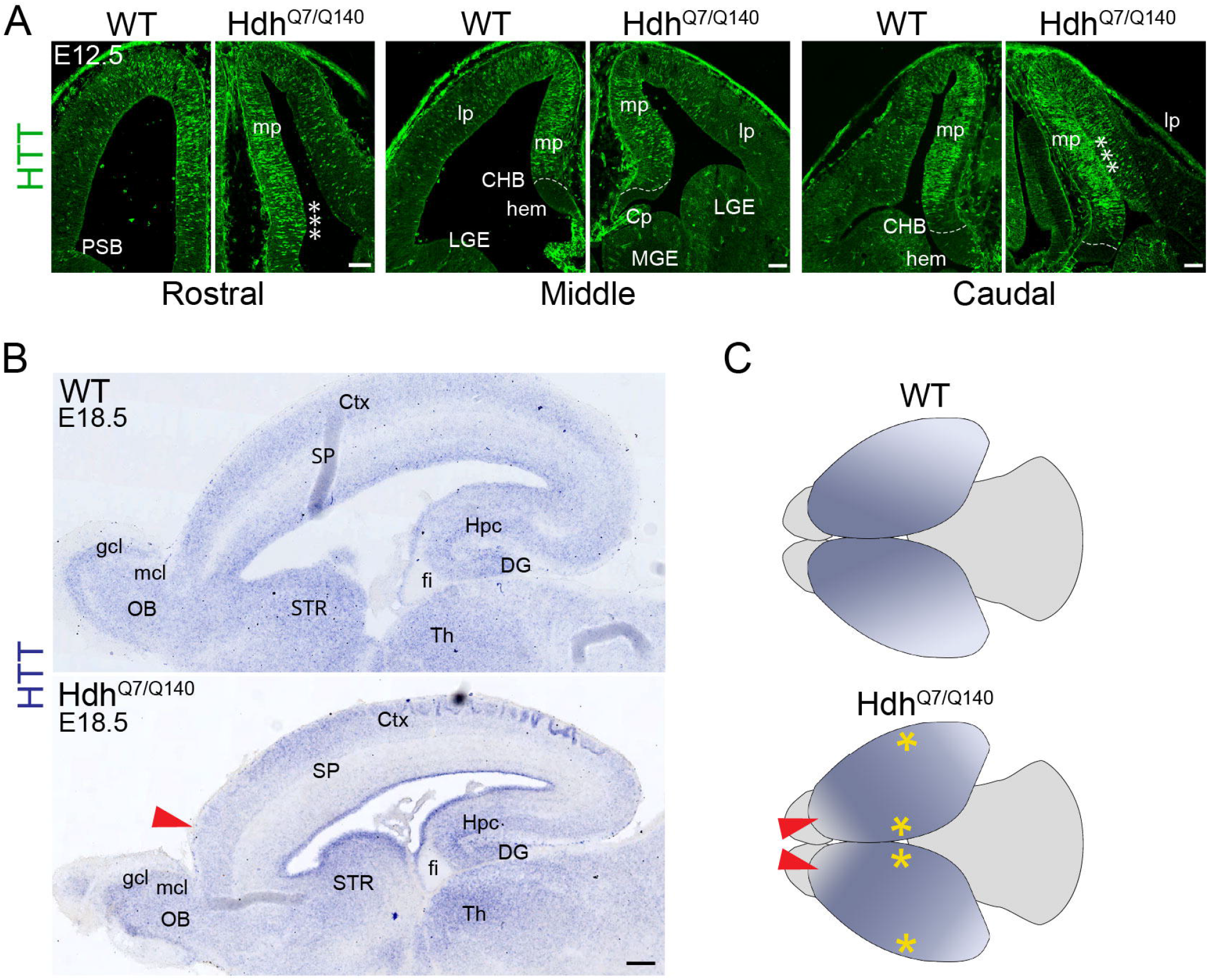
HTT expression in the developing and post-natal mouse brain. (A-B) WT and Hdh^Q7/Q140^ cortical sections were immunostained for HTT (A) and processed for *in situ* hybridization at indicated times (B). (C) Schematic representation of the gradient expression pattern observed in the cortex of WT and Hdh^Q7/Q140^ mice (Fig1. B). The changes of *Htt* expression in HD are indicated by the yellow stars, and the shift of antero-posterior gradient by red arrowheads. Scale bars: 200µm (A) and 300 µm (B). CHB: cortical hem boundary, cp: choroid plexus, Ctx: cortex, hem: cortical hem, DG: dentate gyrus, fi: fimbria; gcl: granular cell layer, hpc: hippocampus, LGE: lateral ganglionic eminence, lp: lateral pallium, mcl: mitral cell layer, MGE: medial ganglionic eminence, mp: medial pallium, OB: olfactory bulb, PSB: pallial subpallial boundary, SP: subplate, Th: thalamus.

We then evaluated *HTT* expression at the end of embryogenesis (E18.5) (Fig. 1B). Within the cortex, *HTT* was mostly detected in the proliferative zone above the lateral ventricle, the upper layers and the thin layer 6b or subplate. In upper layers, the expression was higher in the rostral part of the cortex suggesting an antero-posterior gradient of expression. *HTT* was also expressed in the striatum, the thalamus, the hippocampus and the dentate gyrus. Consistent with the exclusion of *HTT* staining from the cortical hem at E12.5, *HTT* expression was absent from the derivative structure at 18.5, the fimbria of the hippocampus. *HTT* was expressed in the granular and mitral cell layers of the olfactory bulb. In HD brain, the expression of *HTT* was maintained in all quoted structures above, however, in the upper layers the rostral expression appeared downregulated (Fig. 1B). Consistent with the modification of HTT distribution in the medial pallium, the *HTT* gradient was higher in the medial region of the HD sample than in the WT sample (Fig. 1C, yellow stars) and associated to a shift of the anteroposterior gradient (Fig. 1C, red arrows). Furthermore, the expression of mHTT was enriched at the ventricular zone as previously reported^**18**^.

We next determined the size of the hemispheres in control and HD animals. We measured the surface of hemispheres on dorsal views of total head at E12.5 and on dissected brains at E18.5 and P7 (Fig. 2A and 2B) and their length from the medio-frontal tip to the latero-caudal tip (Fig. 2C). The width of hemispheres was measured at 4 levels along antero-posterior axis (Fig. 2D). Overall, all measured parameters were similar at all stages in control and Hdh^Q7/Q140^ mice.

**Figure 2.**
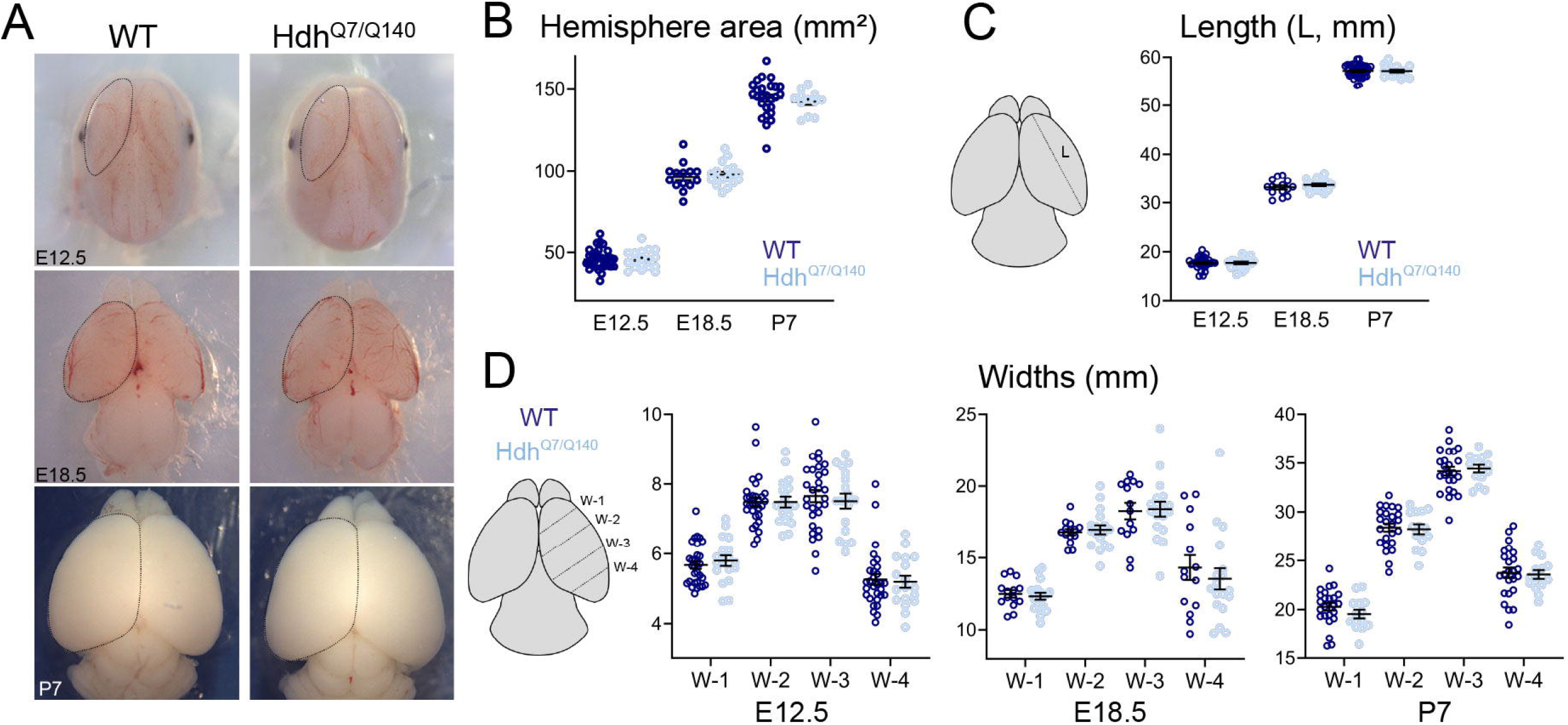
The cerebral hemispheres of Hdh^Q7/Q7^ and Hdh^Q7/Q140^ mice develop at a similar size. (A) Dorsal views of E12.5, E18.5 and P7 WT and Hdh^Q7/Q140^ brains. (B) The graph shows the quantification of dorsal cortical surface areas. (C) The graph shows the measure of the maximal length (L) of the hemispheres between the rostral and caudal apex as indicated in the diagram on the left. (D) Graphs show the measure of four widths (W-1, W-2, W-3, W-4) along the antero-posterior axis as indicated in the diagram on the left. Data represent mean ± SEM. Unpaired t-test (n > 17).

These data indicate that despite alterations of *HTT* expression in Hdh^Q7/Q140^ mice, no severe disorganisation of the brain hemispheres is visible from early development to postnatal stages.

### Diencephalic development is disturbed in Hdh^Q7/Q140^ brain

At the beginning of the second week of embryonic development, the interaction between different signaling pathways in the forebrain prompts the induction of the expression of transcription factors in gradients for the normal development of the cortex^**19– 21**^. This is the case for the telencephalon which dorso-ventral, rostro-caudal and medio-lateral positions are established through the action of several morphogens: sonic hedgehog (*Shh*), in the ventral telencephalon (primordium of hypothalamus), the fibroblast growth factor 8 (*Fgf8*), in the anterior neural ridge, the wingless-type MMTV integration site 3a (*Wnt3a*) and bone morphogenetic protein 4 (*Bmp4*), expressed in the cortical hem and diencephalic territories.

Whole mount *in situ* hybridization of Hdh^Q7/Q140^ embryos at E10.5 revealed that *Shh* expression was similar in control and HD embryos in various territories such as limb buds, the notochord and the zona limitans intrathalamica (Fig. 3A). We next dissected the neural tube of HD embryos and found that *Shh* expression in the anterior hypothalamus was particularly decreased as compared to control animals (Fig. 3A, white asterisk).

**Figure 3.**
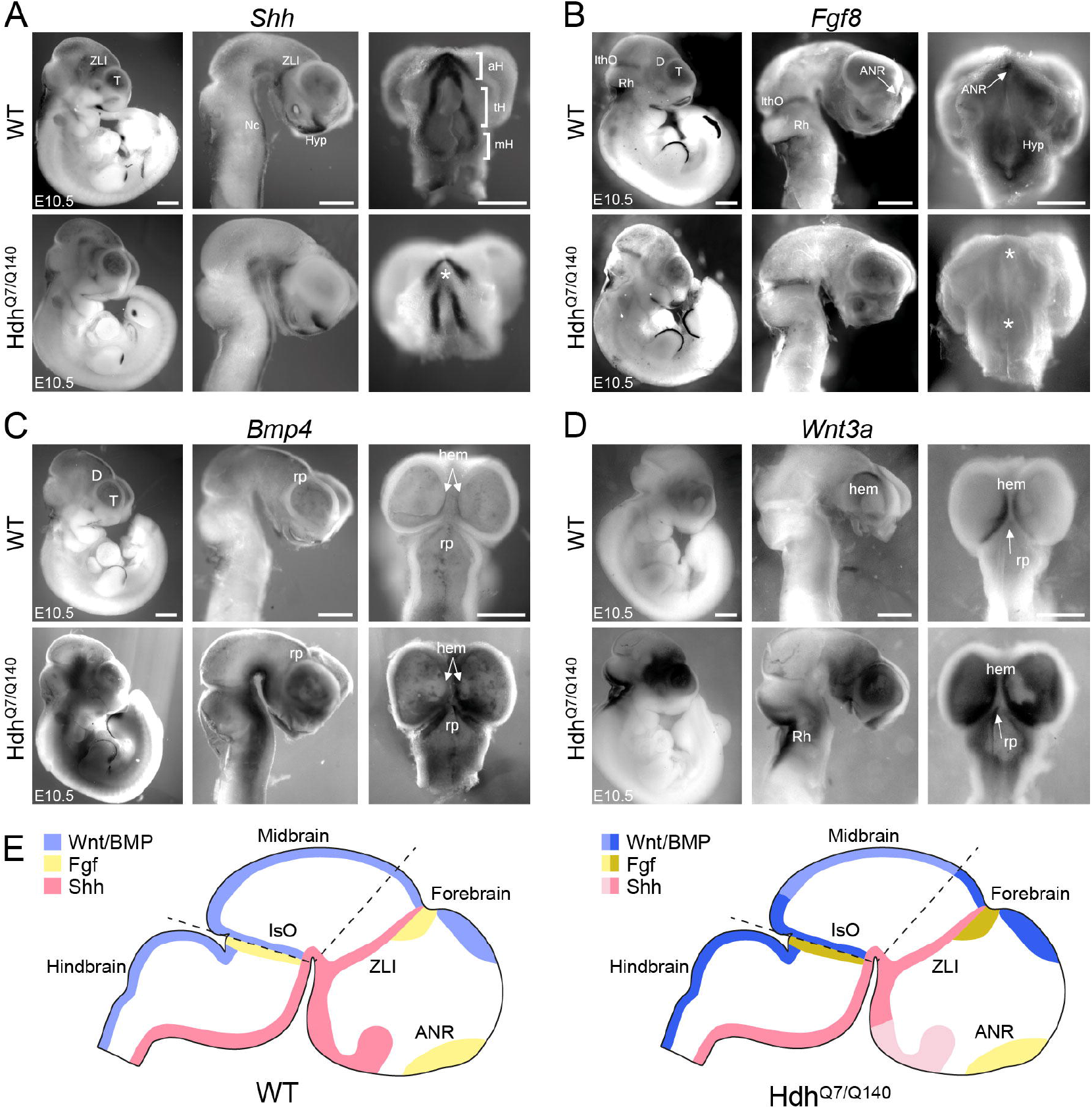
Reduced Shh and Fgf signalling and increased Wnt and Bmp signalling in developing HD brain. (A-D) Whole mount *in situ* hybridization of *Shh* (A), *Fgf8* (B), *Bmp4* (C) and *Wnt3a* (D) in WT and Hdh^Q7/Q140^ mouse embryos at E10.5. From left to right: representative pictures of the whole embryo, the dissected neural tube and a forebrain ventral view for *Shh* and *Fgf8* or a dorsal view for *Bmp4* and *Wnt3a*. Asterisks indicate the reduction of *Shh* and *Fgf8* expression in the Hdh^Q7/Q140^ brains. (E) Schematic representation of changes in the expression of Shh, Fgf and Wnt/BMP signalling in patterning centers. Adapted from Suárez *et al*., 2014^**83**^. Scale bars: 500µm. aH: anterior hypothalamus, ANR: anterior neural ridge, D: diencephalon, hem: cortical hem, Hyp: hypothalamus, ithO: isthmus organizer, mH: mamillary hypothalamus, Nc: notochorde, Rh: rhombencephalon, rp: roof plate, tH: tuberal hypothalamus, T: telencephalon, ZLI: zona limitans intrathalamica.

*Fgf8* expression in the isthmic organizer and dorsal diencephalon was similar in HD and control embryos, but decreased in the anterior neural ridge, hypothalamus and dorsal diencephalon of HD animals (Fig. 3B). Ventral views (Fig. 3B, right panels) also revealed downregulation of *Fgf8* expression in the anterior neural ridge and the primordium of the hypothalamus (Fig. 3B, white asterisks), visible on dissected neural tubes (Fig. 3B, middle panels).

Finally, we addressed the expression of *Bmp4* and *Wnt3a* (Fig. 3C and 3D). In Hdh^Q7/Q140^ brains, *Bmp4* and *Wnt3a* expression was increased in the telencephalic hem and in the diencephalic roof plate which might affect the dorso-ventral patterning of the diencephalon^**22**^.

Thus, the diencephalic expression of signaling molecules and morphogens is modified in E10.5 Hdh^Q7/Q140^ embryos with an overall downregulation of these factors in the ventral part and upregulation in the dorsal part of the forebrain (Fig. 3E) with potential consequences on the organization of the cortical areas.

### Emx2, Lhx2, Sp8, Couptf1 and Pax6 expression is altered in Hdh^Q7/Q140^ mice

Patterning factors such as the ones dysregulated in HD (Fig. 3) subsequently induce the expression of the transcription factors *Emx2, Lhx2, Couptf1* and *Sp8* that execute region-specific neurodevelopmental programs and thus cortical arealization^**23–27**^. We assessed the medio-lateral expression of the genes encoding these transcription factors in HD using *in situ* hybridization at E12.5.

The homeodomain transcription factor EMX2 (Empty Spiracles Homeobox 2) promotes caudomedial fates and is upregulated by Wnt and Bmp^**28**^. LHX2 (LIM homeobox 2) permits the specification of the cortical primordium by suppressing alternative fates corresponding to the hem, antihem, and the paleocortex^**29**^. *Emx2* and *Lhx2* are expressed similarly in a gradient with high caudal to low rostral levels. In HD mice, the gradient pattern of expression of *Emx2* and *Lhx2* was conserved but with an overall downregulation in HD (Fig. 4A and 4B). We also noted a downregulation of *Lhx2* in the proliferative zone of the lateral and medial ganglionic eminences (Fig. 4B).

**Figure 4.**
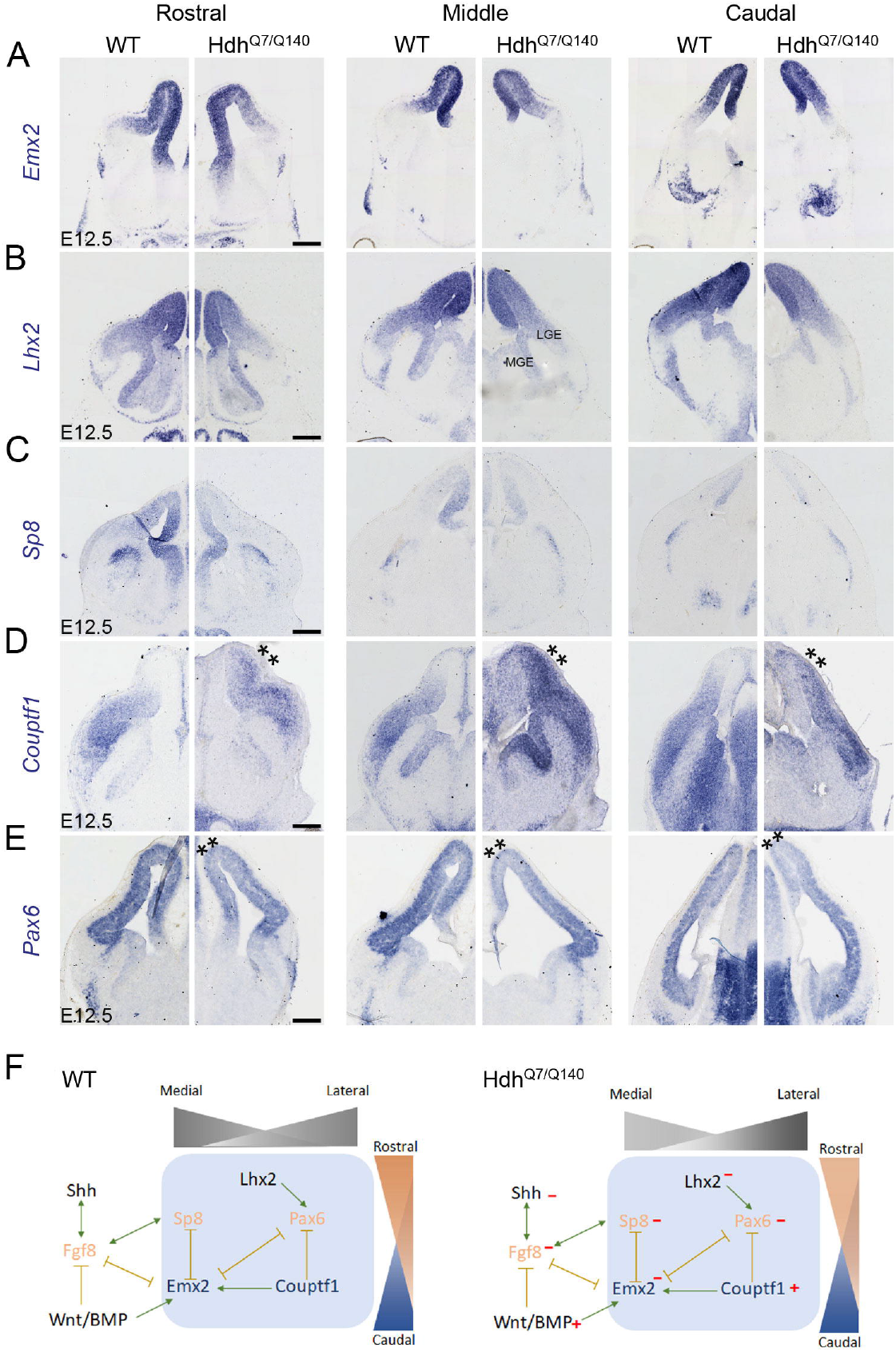
The medio-lateral and rostro-caudal gradients of transcription factors are altered in Hdh^Q7/Q140^ brain. (A-E) Cortical coronal sections of E12.5 mouse brains are processed for *in situ* hybridization for *Emx2* (A), *Lhx2* (B), *Sp8* (C), *Couptf1* (D) and *Pax6* (E) at rostral, mid and caudal levels of sectioning. Asterisks indicate the change of transcription factor expression. (F) Schematic representation of the regulatory network of signalling molecules and transcription factors in control and Hdh^Q7/Q140^ developing brain. Scale bars: 200µm. LGE: lateral ganglionic eminence, MGE: medial ganglionic eminence.

SP8 (Specificity protein 8) promotes frontal/motor areas, whereas COUPTF1 (Chicken ovalbumin upstream promoter) is known to repress them^**30,31**^. *Sp8* is induced by FGF8 and is expressed at the highest levels in the medial and anterior parts of the dorsal telencephalon^**32**^. In accordance with our observation that *Fgf8* expression was decreased in Hdh^Q7/Q140^ embryos, *Sp8* expression was downregulated in the cortical epithelium and medial ganglionic eminences (Fig. 4C). *Couptf1* is expressed in a high-caudolateral to low rostro-medial gradient in the cortex. In the telencephalon of Hdh^Q7/Q140^ embryos, *Couptf1* was expressed in a gradient along both axes as in WT animals. However, its expression was higher and more extended in the caudal part (Fig. 4D). This result is in agreement with the negative regulation between *Sp8* and *Couptf1*^**31**^.

PAX6 (Paired Box 6) specifies rostro-lateral cortical identities and shows a high rostro-lateral to low caudo-medial gradient^**24,27**^. *Pax6* expression was reduced in the dorsal telencephalon of Hdh^Q7/Q140^ embryos (Fig. 4E). Its decrease is consistent with the increased *Couptf1* expression, as these factors act antagonistically during cortical arealization^**29,31**^.

Furthermore, the dorsal limit of these different transcription factors was identical in WT and HD mice, showing that deregulation does not affect the dorso-ventral patterning of the telencephalon. To confirm this, we examined the expression of *Ascl1*, a marker of the ventral telencephalon with a restricted subpallial expression pattern. The expression was similar in control and HD samples, indicating that the telencephalon does not become ventralized in HD (Fig. S1).

Together, these findings indicate that mHTT disrupts the expression of key transcription factors controlling cortical arealization without altering dorsoventral telencephalic identity (Fig. 4F). Although global gradient organization is largely preserved, the imbalance between rostro-lateral and caudal patterning programs suggests an early defect in cortical regionalization in HD.

### mHTT modulates the area patterning of cerebral cortex

We next assessed whether the deregulation of transcription factors involved in cortical patterning during the embryonic stage leads to alterations in cortical arealization in HD. Whole-mount *in situ* hybridization was performed on postnatal day 7 (P7) brains, a stage at which neurons have reached their laminar positions and cortical area boundaries can be identified based on gene expression patterns^**33**^. In control animals, *Ror*β (Rare-related orphan receptor β) expression delineated the visual (V1), primary somatosensory (S1), and auditory (A1) cortices, with weaker expression detected in some higher-order visual areas^**34**^ (Fig. 5A). *Lmo4* (LIM-domain-only 4) was expressed rostrally in the frontal/motor cortex (M1) and medially in the cingulate cortex, while being excluded from primary sensory areas^**35**^ (Fig. 5B). *Cad8* (Cadherin-8) predominantly labeled M1 and V1^**15**^ (Fig. 5C).

**Figure 5.**
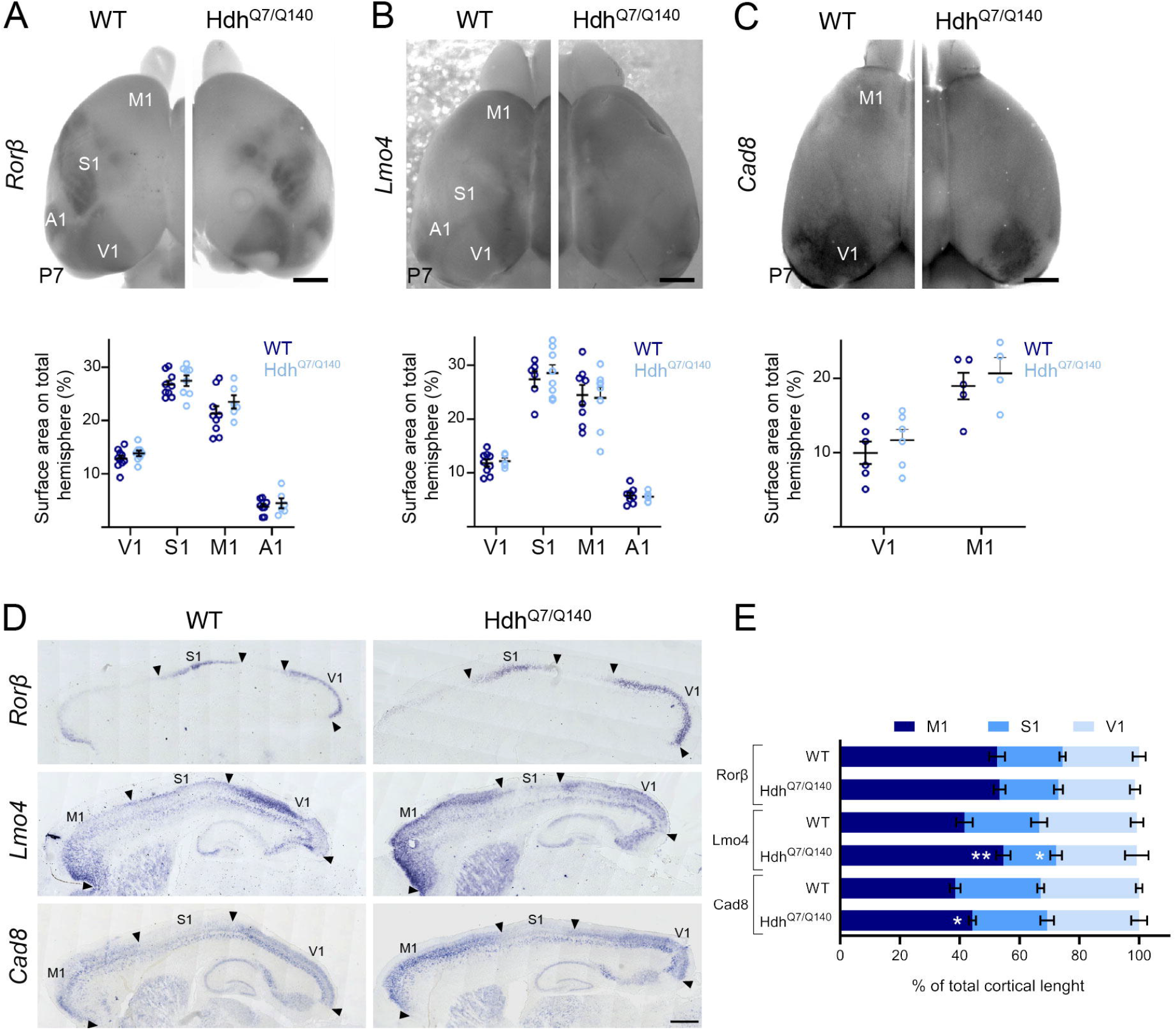
HTT modulates the patterning of neocortical areas. (A-C) Dorsal views of WT and Hdh^Q7/Q140^ P7 brains processed for whole mount *in situ* hybridization for *Rorβ* (A), *Lmo4* (B) and *Cad8* (C). Graphs show the ratio of the size of each area to that of the entire hemisphere size. (D) Sagittal brain sections of *in situ* hybridization for *Rorβ, Lmo4* and *Cad8* expression. Arrowheads indicate limits of different areas used for the quantifications of area lengths in the graph. (E) Quantification of the motor, somatosensory and visual areas in WT and Hdh^Q7/Q140^ mice. Data are represented as means ± SEM; T-test *p<0.05; **p<0.01. Scale bars: 1mm (A, B and C), 500µm (D). A1: auditory area, M1: fronto/motor area, S1: somatosensory area, V1: visual area.

The total cortical surface area at P7 was unchanged in Hdh^Q7/Q140^ mice compared with controls (Fig. 2B). Quantification of the surface areas of V1, S1, A1, and M1, as defined by the expression of *Rorβ, Lmo4*, and *Cad8*, revealed no abnormalities in the extent of these territories in HD brains (Fig. 5A-C).

We then examined the rostro-caudal positioning of primary cortical areas. *In situ* hybridization for *Rorβ, Lmo4*, and *Cad8* was performed on sagittal sections at P7, and the length of each domain was normalized to the total cortical length (Fig. 5D). *Rorβ* staining showed comparable rostro-caudal extents of S1 and V1 between genotypes. In contrast, *Lmo4* and *Cad8* expression revealed a rostral expansion of M1 in Hdh^Q7/Q140^ brains relative to controls, accompanied by a relative reduction of S1 and V1 territories (Fig. 5E).

Altogether, these data indicate a rostro-caudal shift of primary cortical areas at the surface of the hemispheres in Hdh^Q7/Q140^ mice at P7, suggesting an early mispositioning of motor and sensory domains that could affect the subsequent organization of cortical circuits.

### Hdh^Q7/Q140^ pups exhibit defects in sensorimotor behavior

We thus investigated whether the altered cortical area patterning up to P7 in the motor and somatosensory regions translated into behavioral outcomes during the first postnatal week. We did not observe significant differences in body weight between genotypes at any postnatal day (Fig. 6A). Neurodevelopmental reflexes, also known as primitive reflexes, are commonly used to assess brain development in neonatal pups. The limb grasp reflex relies on spinal reflexes and corticospinal inhibition^**36**^. Interestingly, Hdh^Q7/Q140^ male pups performed better on this test than control males at post-natal day 1 (P1) (Fig. 6B). However, only half of the Hdh^Q7/Q140^ females were able to grasp at P1, compared to nearly 75% of control females. By P2, this difference was reversed in males, with control males outperforming Hdh^Q7/Q140^ males.

**Figure 6.**
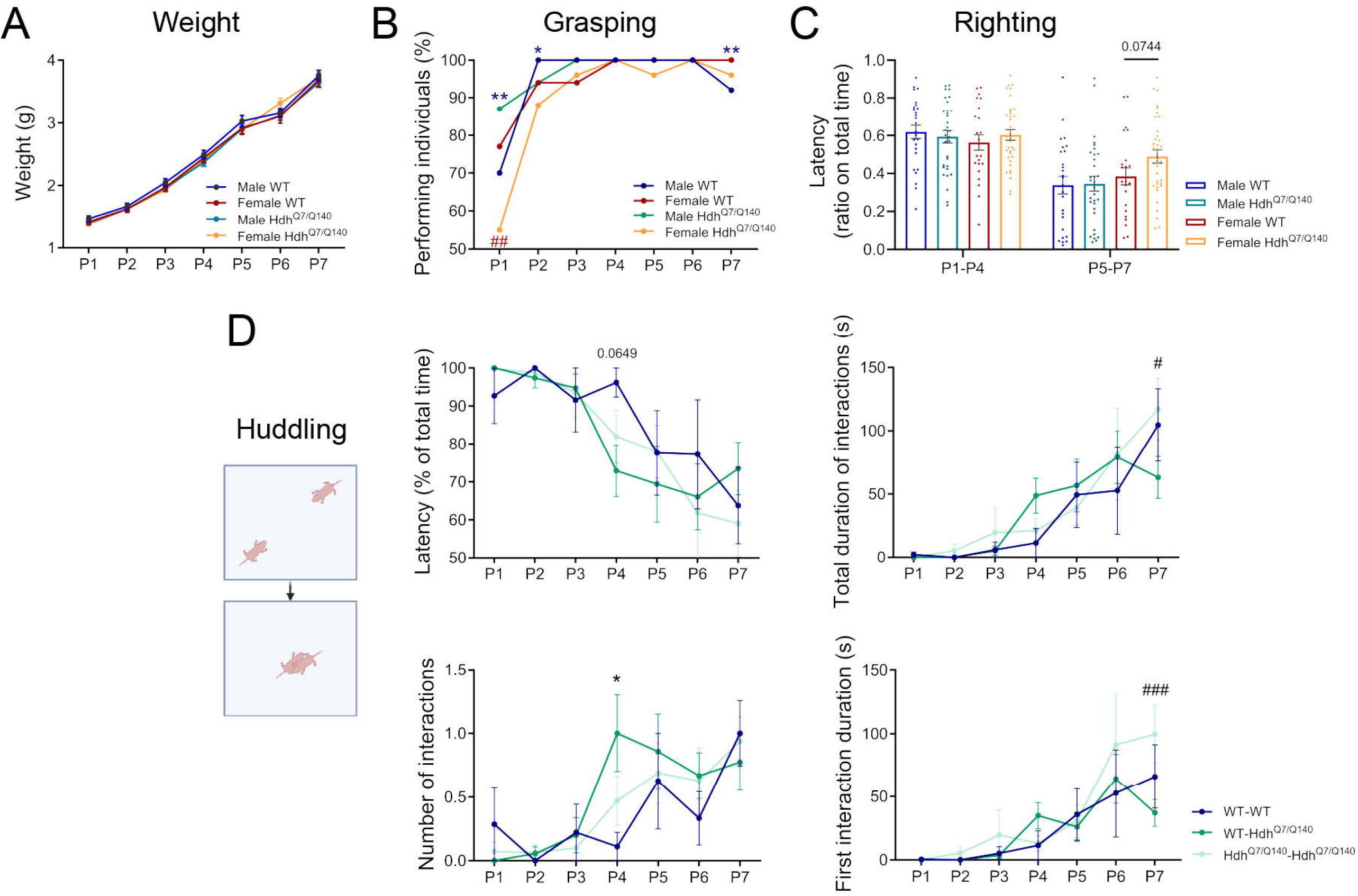
Hdh^Q7/Q140^ pups exhibit defects in sensorimotor behaviour. (A) Body weight monitored from P0 to P7 in WT and Hdh^Q7/Q140^. Two-way ANOVA followed by Tukey’s multiple comparison test (n = 15-36). Grasping (B) and righting (C) behaviours were evaluated from P1 to P7. Data are represented as means ± SEM; Multiple Mann-Whitney tests and Contingency Fisher’s test respectively (n = 23-36) *p<0.05. **p<0.01 comparing male groups, ##p<0.01 comparing female groups. (D) Huddling behaviour is tracked on three pairs of pups: WT-WT, WT-Hdh^Q7/Q140^ and Hdh^Q7/Q140^ Hdh^Q7/Q140^ from P1 to P7. The latency, the total duration, the number of interactions and the first interaction duration are evaluated. Data are represented as means ± SEM; Two-way ANOVA followed by Tukey’s multiple comparison test (n = 6-22), * or #p<0.05, ###p<0.001 Sidak’s comparisons after significant interaction.

We also assessed the righting reflex, which involves vestibular sensation, proprioception, touch sensation, and motor coordination^**16**^. We recorded the time required for pups to roll over and place all four paws on the surface (Fig. 6C). The results were grouped into two stages based on noticeable developmental gains. While no significant differences were observed between sex and genotype from P1 to P4, Hdh^Q7/Q140^ females tended to take a longer time to right themselves at P5–P7 compared to their WT counterparts.

Following cortical maturation, huddling is one of the earliest behaviors associated with the sensorimotor cortex, where pups spontaneously interact for thermoregulation^**37**^. From P1 to P7, pups were separated from their mother and video-recorded in an empty arena in same-sex pairs, with the genotype blinded to the observer (Fig. 6D). The time spent moving until reaching each other was monitored (latency). No specific phenotype was observed from P1 to P3. However, at P4, pairs composed of one WT and one Hdh^Q7/Q140^ pup tended to huddle significantly faster than WT-only pairs. At this stage, these mixed pairs also engaged in more interactions. This suggests that heterogenic pairs recognize each other more quickly, despite their interactions being less stable. This defect persisted at P7, where heterogenic pairs exhibited shorter first interaction compared to homogenic pairs. In contrast, homogenic pairs spent more time interacting than heterogenic pairs.

These findings indicate early impairments when primitive reflexes, motor and sensory functions develop, followed by deficits in huddling behavior that develops later, at the beginning of the second postnatal week.

## DISCUSSION

Our study demonstrates that mHTT alters the expression of signalling molecules and transcription factors essential for cortical areas patterning. In HD, these abnormalities are correlated to abnormal motor and somatosensory behavior during the first postnatal week.

We showed that HTT is expressed in cortical apical progenitors in a high rostro-medial-to-low caudo-lateral gradient and is excluded from the cortical hem and its derivative, the fimbria^**38**^. The cortical hem is a key signalling centre secreting Wnt/BMP molecules^**39**^ required for the development of caudo-medial cortex, hippocampus, and choroid plexus^**40–42**^. Moreover, the cortical hem generates Cajal-Retzius cells that influence the telencephalic development^**43,44**^. HD mice show defects in adult hippocampal neurogenesis and in the organization of the dentate gyrus^**45–50**^. Altered spatial memory in HD patients has also been associated to hippocampal abnormalities^**51–54**^. Consistently, we observed overexpression of *Wnt3a*, the earliest of the Wnt family member selectively expressed in the cortical hem^**41**^. Dysregulated hem-derived Wnt signalling, which controls dentate neuroepithelium proliferation and glial scaffold formation^**44**^, may therefore contribute to early HD-related defects.

Our data suggests an interaction between Wnt and HTT at the cortical hem boundary which is supported by previous studies showing modifications of Wnt pathways in HD^**55**^. Wnt inhibition prevents angiogenic deficits and blood brain barrier impairment in HD iPSC-derived brain microvascular endothelial cells^**56**^. However, the role of mHTT on the Wnt pathway which can act in a canonical or noncanonical way in the cortical hem remains unknow and requires further investigations. It may involve β-catenin which binding to HTT is impaired in primary striatal neurons from HD mice and iPSCs from HD patients^**57,58**^.

Knowledge about neural tube formation and brain morphogenesis in HD remains fragmented. Due to the embryonic lethality of *HTT* knock-out mice, hypomorphic models have been used to describe defects in the epiblast of primitive streak^**59,60**^. *HTT* knockdown in zebrafish embryo revealed defects in the most anterior regions of the neural plate, in the midbrain roofplate and rhombomeric boundaries^**61,62**^. Our observation of *Wnt3a* overexpression in the rhombencephalon is thus consistent with previous studies showing the expansion of Wnt gene expression when *Fgf8* is downregulated^**63,64**^.

The hypothalamus is a ventral forebrain region regulating homeostasis and reproduction; alterations of hypothalamus development can result in endocrine and metabolic diseases. The embryonic hypothalamus forms the rostral-ventral part of the diencephalon. *Shh*, is expressed early in the development in the prechordal plate, the notochord and by the ventral midline of the neural tube. In the diencephalon, *Shh* also has a specific expression pattern in the zona limitans intrathalamica and the rostral-ventral diencephalon (hypothalamus domain) which influences the formation of the diencephalon-telencephalon junction^**65**^. In HD patients, several endocrine alterations arise as diabetes, glucose homeostasis and neuropathological changes in the hypothalamus occur before clinical diagnosis^**66,67**^. In mouse models of HD, studies showed the contribution of gender to HD progression and severity: deregulations of HTT expression in the hypothalamus of female R6/2 and BACHD elicit different effects on metabolic phenotype and their littermates^**68–72**^. In HD, we found Shh-dependent developmental abnormalities in regions at the origin of the future hypothalamus. Consistently, Hippi (a molecular partner of the HTT-interacting protein HIP-1) knockout mice exhibit downregulation of the Shh signaling pathway, along with defects in nodal cilia assembly, leading to developmental abnormalities and characteristic features of impaired left–right axis patterning^**73**^. Whether the HD adult hypothalamic disturbances find their roots in development and the dysregulation of Shh pathway needs further investigations.

We found a downregulation of transcription factors in rostral and medial parts of the developing brain with modification of area regionalization at later stages. The Wnt and BMP pathways induce *Emx2* expression^**74**^. In our model, the cortical hem structure remains intact and *Emx2* is downregulated. In absence of LHX2, different studies showed an expansion of the cortical hem and antihem, and an increase of Wnt/BMP signaling molecules^**29,44,75–77**^. To explain our data, we could hypothesize that transcription factors as FOXG1^**78**^, DMRT5 or DMRT3^**11**^ that interact with Emx2 are also modified in HD.

The cortical patterning is also regulated by COUPTF1, through the repression of the MAPK/ERK signalling, which is altered in HD^**79,80**^. Moreover, COUPTF1 acts on *Pax6* level^**81**^. In the conditional deletion of *Pax6* in cortical progenitors, the hemisphere size is halves and the size of the sensory motor cortex diminishes^**82**^. Therefore, the effect of HD on the position of the fronto-motor area in the neocortical map seems likely to reflect an excess of *Couptf1* in the dorsal brain. *Couptf1* antero-posterior gradient pattern is conserved but we observed a strong overexpression in dorsal and lateral pallium in Hdh^Q7/Q140^. The overexpression of *Couptf1* should trigger an expansion of caudal cortical areas and concomitant expansion of rostral areas^**23,30**^. However, the downstream signaling of COUPTF1 in the context of HD remains to be elucidated.

The cortical layer formation and area patterning are coordinated processes to ensure the proper cytoarchitecture and establishment of cognitive and social behaviours. Previous studies revealed roles of HTT at different steps of the cerebral cortex development which are all impacted in HD^**3–6,55**^. We showed that an early transient treatment during the first week of life of HD mice with ampakine (an agonist of AMPA receptors) restore the dendritic arborization and sensorimotor function in HD pups^**10**^. Our observations here that sensorimotor defects occur in Hdh^Q7/Q140^ pups in the specific windows of P4-P6 days match with the pic of synaptogenesis in the cerebral cortex and hippocampus^**37**^. We thus further consolidate the idea of an early transient period of changes that are compensated later but are of interest for therapeutic correction pre-empting the HD brains own compensation.

---

**Figure S1.**
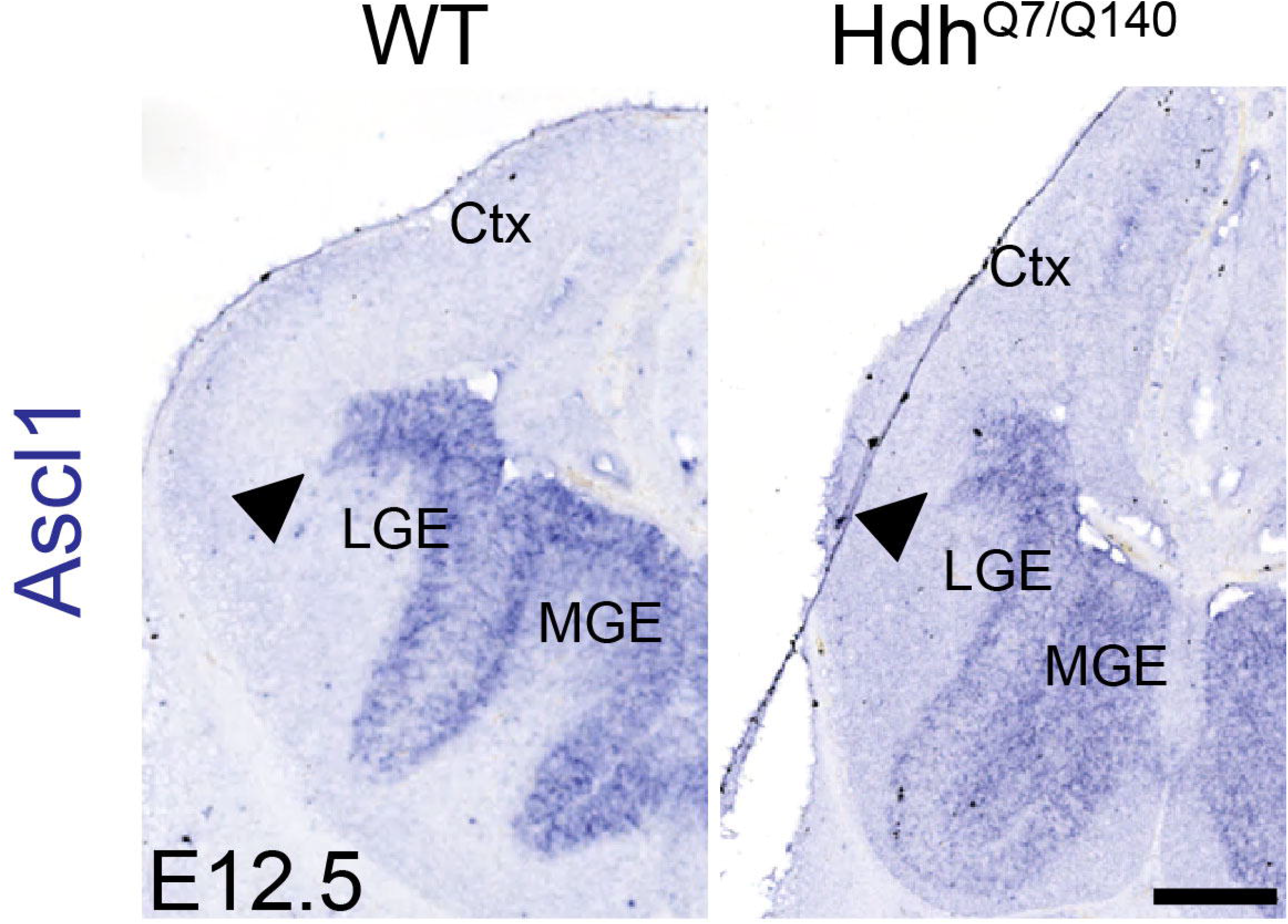
The expression of the ventral determinant Ascl1 is similar in WT and HdH^Q7/Q140^ brains. *In situ* hybridization of E12 embryos. *Ascl1* expression is conserved in the ganglionic eminence in HD and the dorso-ventral limit is visible at the pallium-subpallium boundary (indicated by the arrowheads). Scale bar: 200µm. Ctx: cortex, LGE: lateral ganglionic eminence, MGE: medial ganglionic eminence.

## Acknowledgements

We are grateful to the staff of the Photonic Imaging Center of Grenoble Institute of Neuroscience (PIC-GIN, University Grenoble Alpes – INSERM U1216), which is part of the ISdV core facility and certified by the IBiSA label; L.C. Diaz and J. Vollaire (Optimal platform, Institute for Advanced Biosciences, Grenoble, France) for technical help. We thank F. Agasse for helpful comments and suggestions on the manuscript.

## Ethical considerations

There are no human participants in this article and informed consent is not required. All experimental procedures were performed in authorized establishments (Grenoble Institute of Neurosciences, license #B3851610008) in strict accordance with the local animal welfare committee (Comité Local Grenoble Institute Neurosciences, C1EA-04), EU guidelines (directive 2010/63/EU) and the French National Committee (2010/63) for care and use of laboratory animals under the supervision of authorized investigators (permission APAFIS#32445-2021062417002297 v3).

## Authors contributions

CL: Data curation, Formal analysis, Investigation, Methodology, Validation, Visualization, Writing – original draft, Writing – review & editing. LR: Conceptualization, Data curation, Formal analysis, Methodology, Project administration, Supervision, Visualization, Writing – original draft, Writing – review & editing. SH: Funding acquisition, Project administration, Resources, Supervision, Visualization, Writing – review & editing.

